# Functional stability despite high taxonomic turnover characterizes the Ulva microbiome across a 2,000 km salinity gradient

**DOI:** 10.1101/2024.06.20.599874

**Authors:** Luna M. van der Loos, Sophie Steinhagen, Willem Stock, Florian Weinberger, Sofie D’hondt, Anne Willems, Olivier De Clerck

**Author notes:** corresponding author: Luna van der Loos.

## Abstract

The green seaweed Ulva depends on its associated bacteria for morphogenesis and is an important model to study algal-bacterial interactions. Ulva-associated bacteria exhibit high turnover across environmental gradients, leading to the hypothesis that bacteria contribute to the acclimation potential of the host. Yet little is known about the variation in the functional profile of Ulva-associated bacteria in relation to environmental changes. To test which microbial functions shift alongside a strong environmental gradient, we analysed microbial communities of 91 Ulva samples across a 2,000 km Atlantic–Baltic Sea salinity gradient using metagenomic sequencing. Metabolic reconstruction of 639 metagenome-assembled genomes revealed widespread potential for carbon, sulphur, nitrogen, and vitamin metabolism, including amino acid and vitamin B biosynthesis. While salinity explained 70% of taxonomic variation, it only accounted for 17% of functional variation, indicating extensive functional stability. The limited variation was attributed to typical high-salinity bacteria exhibiting enrichment in genes for thiamine, pyridoxal, and betaine biosynthesis. These metabolic modules likely contribute to oxidative stress mitigation, cellular osmotic homeostasis, and membrane stabilization in response to salinity variations. Our results emphasise the importance of functional profiling to understand the seaweed holobiont and its collective response to environmental change.

## Introduction

Microbial communities interact with eukaryotic hosts in every ecosystem, yet our understanding of how (variation in) microbial metabolisms impact host performance is very limited. Seaweeds, or macroalgae, are a group of important coastal inhabitants. As ecosystem engineers and primary producers, they are an essential source of food and they provide shelter and habitat for diverse marine organisms ^1^. Health, reproduction, and development of the macroalgal host — and indirectly their ecosystem functioning — are strongly dependent on the associated symbionts. Beyond the exchange of key nutrients, vitamins, and secondary metabolites, the microbial biofilm forms a physical and chemical barrier acting as a “second skin” that protects the host and modulates its interactions with the environment ^2–4^. Beneficial bacteria can, for example, shield the host against pathogens ^5^, stimulate algal growth ^6^, or mitigate the adverse effects of environmental pollution ^7^.

Species of the green seaweed Ulva are particularly well-studied and are considered a model system to study algal-bacterial interactions ^8^. The relation between Ulva and its symbionts, together often referred to as a holobiont, is so indispensable that the seaweed fails to develop its typical leaf- or tube-like morphology in the absence of particular bacteria ^9,10^. Morphological development is induced by chemical compounds produced by bacteria that trigger cell wall development, rhizoid formation, and cell division ^11^. Initial studies identified a combination of two complementary strains (a Roseovarius and a Maribacter strain) that were necessary for full morphogenesis in Ulva mutabilis. Since then, multiple strains have been identified that have the same capacity ^12^. This functional redundancy across taxa is important for the host in relation to shifts in bacterial communities.

Seaweed-associated bacterial communities are far from static. Although a small core community can sometimes be identified, the bacteria form a highly dynamic community that fluctuates through time and space ^13,14^. Abiotic factors, such as temperature, light, and salinity influence the community composition, as does the host itself ^15–17^. Prior research has mainly focused on dynamics in the taxonomic composition of the communities rather than the functional potential that the community harbours. Taxonomic profiling is based on 16S rRNA amplicon sequencing that in recent years has become widely accessible due to increased speed and reduced costs. Genome-wide studies or metagenomic sequencing that give information on the functional potential of the bacterial communities in comparison are more expensive and the sample size in those studies is often limited. Metagenomic functional profiling in, for example, Sargassum spp. ^18,19^, Pyropia haitanensis ^20^, and Nereocystis luetkeana ^21^ highlighted the importance of the bacterial biofilm in terms of vitamin production, polysaccharide degradation, and nutrient cycling. However, these studies were based on a relatively small number of replicates (between 3–7 samples) and therefore do not capture differences across environments. To understand how the functional potential of a bacterial community responds to environmental change and to evaluate the potential consequences for the seaweed host, it is essential to expand functional profiling on a broader scale across diverse environmental gradients ^22^.

Ulva species are known for their broad salinity tolerance, thriving in environments ranging from freshwater habitats to highly saline conditions. The Baltic Sea is a particularly interesting area to study Ulva, as it hosts more than 15 species distributed across its distinctive salinity gradient ^23^. This relatively stable gradient stretches across more than 2,000 km of distance, transitioning from near freshwater conditions in the innermost parts towards fully marine conditions on the North Sea side ^24^. While the distribution of certain Ulva species is confined to higher salinity levels, other species like U. intestinalis and U. linza can be found throughout the entire range. Previous studies showed that salinity also strongly structures the taxonomic composition of Ulva- associated bacterial communities ^25^. The question remains, however, whether these taxonomic shifts in response to salinity are reflected by corresponding changes in functional profile of the same bacterial communities, or if functional traits are conserved across the environmental gradient ^26^.

To address these knowledge gaps and enhance our understanding of the Ulva holobiont model system, we investigated the taxonomic and functional composition of Ulva- associated bacteria across the Baltic–Atlantic salinity gradient using metagenomic sequencing. The aim of this study was twofold: (i) to provide a comprehensive overview of the functional potential within the Ulva bacteriome based on a large number of samples (n=91), and (ii) to assess if the functional potential of the microbial community was stable across an environmental gradient. We expected that the functional repertoire of Ulva-associated bacteria would be rich in carbon utilisation and nutrient cycling pathways, as these functions are traditionally associated with seaweed microbiomes ^20,21^. Given that salinity has been identified as a major driving force of global bacterial diversity and community structure ^27^, we also hypothesized pronounced changes in the functional profile of the Ulva-associated bacterial community, particularly in functions related to osmoprotection.

## Results and discussion

### Ulva-associated bacteria utilize host-derived carbon and sulphur

Rich in organic carbon, oxygen, and nutrients, the algal surface provides an ideal habitat for the growth of microbes ^28,29^. The microbial community reciprocates by supplying the algal host with phosphate, nitrogen, and vitamins in return ^2,30^. Such an exchange of key nutrients is an essential feature of the Ulva holobiont. We used metagenomic sequencing to comprehensively describe the functional profile of Ulva-associated bacteria, obtaining a large dataset of metagenome-assembled genomes (MAGs) that were annotated with the Carbohydrate-Active EnZymes database (CAZy) ^31^ and Kyoto Encyclopedia of Genes and Genomes (KEGG) ^32,33^. Across 91 Ulva microbiome samples (Table S1) and 639 MAGs (Table S2), we identified a total of 6,525 KO (KEGG Orthology) terms and 399 KEGG modules (Table S3). MAGs with an estimated completeness of >90% contained on average 59 KEGG modules (ranging from 33-94 modules). Apart from basic cellular metabolic pathways (e.g., fatty acid biosynthesis, RNA polymerases, DNA polymerases, ribosome, F-type ATPases, cytochrome C oxidase, etc.), the Ulva-associated bacterial communities contained a range of functions that are clearly linked to the association with its host. This included the widespread potential of bacteria to utilize host-derived carbon and sulphur.

All 639 MAGs were able to utilize organic carbon as their primary energy source through, e.g., complex carbon degradation, glycolysis (KEGG modules M00001+M00002), and the tricarboxylic acid (TCA) cycle (M00009) (Fig. 1a). Likewise, many bacteria were capable of utilizing sulphur compounds produced by the host. Sulphur metabolism related genes in Ulva-associated communities were predominantly involved with assimilatory sulphate reduction (M00176), leading to the formation of sulphite and, ultimately, sulphide (Fig. 1a). In addition, 137 MAGs were able to oxidize thiosulphate (the result of the incomplete oxidation of sulphides) through sulphur-oxidizing proteins SoxAB and SoxXYZ that together form the Sox complex (M00595) (Fig. 1a). Due to the presence of sulphate esters in cell wall polysaccharides, Ulva species have a relatively high sulphur content ^34,35^ and our findings support the hypothesis that the widespread prevalence of sulphonates in marine algae contributes to the high abundance of sulphonate-degrading bacteria in marine habitats ^36^.

**Fig. 1.**
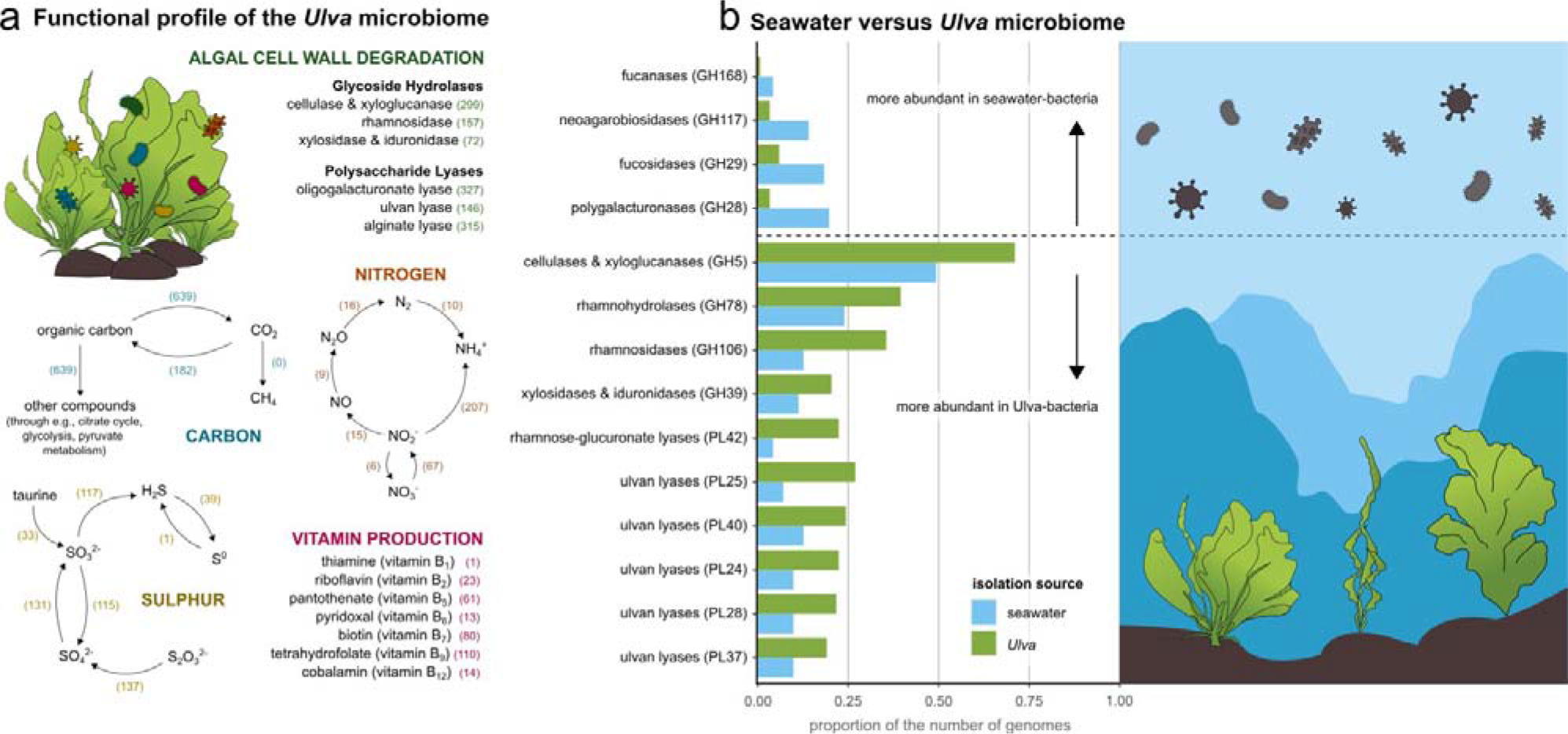
Functional profile of the Ulva microbiome. (a) Overview of the metabolic potential of the Ulva microbiome, highlighting vitamin production (purple), algal cell wall degradation (green), and the metabolism of carbon (blue), nitrogen (orange), and sulphur (yellow). The number of identified metagenomic assembled genomes (MAGs) that are capable performing the reactions are displayed in brackets. (b) The proportion of the number of seawater-isolated (blue) or Ulva-isolated (green) bacterial genomes than contained a specific CAZY family (polysaccharide lyases = PL, glycoside hydrolases = GH).

Other prevalent sulphur cycling enzymes in Ulva-associated bacteria were related to dimethylsulfoniopropionate (DMSP) metabolism, an organosulphur compound produced by Ulva ^37,38^. DMSP has a wide range of ecophysiological functions and in the model species Ulva mutabilis it is known to mediate the interaction with the symbiont Roseovarius sp. MS2. This symbiont releases morphogenetic compounds that stimulate algal morphogenesis and growth ^39^. Interestingly, although the bacterial strain is attracted by the release of DMSP and rapidly takes up the compound, its growth is not affected by DMSP. Instead, DMSP is converted into dimethylsulphide (DMS) and methanethiol (MeSH). In our dataset, the capacity to convert DMSP to DMS was almost exclusively restricted to a few genera within the Rhizobiaceae (the undescribed genus JAALLB01) and Rhodobacteraceae (predominantly Jannaschia, Roseovarius, Sulfitobacter, and Tateyamaria). This observation aligns with previous findings that another well-known morphogenesis-inducing symbiont of Ulva, Maribacter sp. MS6 (belonging to the Flavobacteriaceae), is not attracted by DMSP ^39^.

Apart from examining carbon and sulphur metabolic functions organized in KEGG modules, we also screened the Ulva-associated bacterial MAGs for the presence of carbohydrate-active enzymes using the CAZy database. The CAZy database categorizes enzymes involved in degrading, modifying, or creating glycosidic bonds into families, including glycoside hydrolases (GHs) and polysaccharide lyases (PLs). A total of 121 different GH families and 26 different PL families were identified within Ulva-associated MAGs. Our results demonstrated that a substantial majority [97%] of the bacteria in Ulva-associated communities are capable of breaking down complex polysaccharides constituting the Ulva cell wall. The Ulva cell wall is mainly composed of cellulose, glucuronan, ulvan, and xyloglucan, which together make up 45% of the dry-weight biomass ^40^. Ulvan, a key component, is composed of sulphated rhamnose, xylose, and uronic acids (D-glucuronate and iduronic acid) ^40^. The initial steps in the breakdown of ulvan can be catalysed by ulvan lyases (sulfatases) ^41^, but additional glycoside hydrolases are needed to obtain monomeric sugars ^42^. Ulvan lyases [amongst others CAZy family PL24, PL25, and PL37] were found in 146 MAGs (Fig. 1a), many of which were typical Ulva-associated bacteria, such as Lewinella, Nonlabens, Algibacter, Glaciecola, and Polaribacter ^12^. Prevalent glycoside hydrolase families included GH3 [553 MAGs], GH16 [305 MAGs], and GH5 [299 MAGs]. Family GH3 includes, amongst others, xyloglucan-specific exo-β-1,4-glucosidase (EC 3.2.1.-), xylan β-1,4-xylosidase (EC 3.2.1.37), several glucanases, and several other glucosidases. Enzymes of family GH16 are active on β-1,4 or β-1,3 glycosidic bonds in various glucans and galactans and includes xyloglucan:xyloglucosyl transferase (EC 2.4.1.207). Members of the GH5 family are mainly active on cellulose, but this family also includes high specificity xyloglucanases.

### Nitrogen metabolism and amino acid biosynthesis

Unlike animals, seaweeds can synthesise essential amino acids themselves and relatively little is known about the uptake of dissolved organic nitrogen compounds by seaweeds ^43^. It is likely, however, that seaweeds and bacteria do exchange amino acids, especially when dissolved inorganic nitrogen availability is low ^44^. The biosynthesis of multiple amino acids was part of the core modules of the Ulva-associated bacteria (with 0.65- 1.42% relative abundance in at least 99% of samples). This included the biosynthesis of proline (KEGG module M00015), threonine (M00018), lysine (M00016, M00526, M00527), valine/isoleucine (M00019), serine (M00020), cysteine (M00021), tryptophan (M00023), histidine (M00026), ornithine (M00028), leucine (M00432), and arginine (M00844). Of these core modules, cysteine biosynthesis was detected in the fewest number of MAGs (222 MAGs), while histidine biosynthesis was found in the highest number of MAGs (443 MAGs). Likewise, genes necessary for glycine biosynthesis (from threonine or serine), alanine biosynthesis (from pyruvate) and glutamine biosynthesis (from glutamate), as well as the conversion between aspartate and asparagine were prevalent in the majority of the MAGs.

Nitrogen metabolism potential largely centred on the conversion from nitrate and nitrite to ammonia (Fig. 1a). A large number of 174 MAGs contained the nrfAH or nirBD genes that are essential for nitrite reduction to ammonia, which is the final step in dissimilatory nitrate reduction to ammonia. A total of 30 MAGs exhibited the genes for the full dissimilatory nitrate reduction to ammonia, utilizing nitrate respiration for energy production (M00530). Only seven MAGs, all belonging to the Cyanobacteriia, demonstrated complete assimilatory nitrate reduction capability, involving the reduction of nitrate to ammonia for biosynthesis purposes (M00531). Nevertheless, the first step in this process (nitrate to nitrite reduction mediated by the narB gene) was identified in 53 MAGs, and the subsequent step (nitrite to ammonia reduction facilitated by the nirA gene) was detected in 45 MAGs. The narB gene was predominantly observed in Flavobacteriaceae and Saprospiraceae, while the nirA gene was prevalent in Akkermansiaceae and Pirellulaceae. Two MAGs harboured all the genes necessary for the denitrification process involving the reduction of nitrate and nitrite to nitrogen (M00529), while none of the MAGs carried genes associated with nitrification (the oxidation of ammonia to nitrate; M00528). Finally, ten MAGs from diverse families showed the capacity to fix nitrogen (M00175).

Previous studies demonstrated that observed amino acid uptake rates in Ulva were highest for glycine and alanine and that while Ulva prefers inorganic nitrogen, the organic compounds may also play a significant role ^45,46^. A transcriptomic study in Laurencia dendroidea showed that the associated bacteria had a higher relative contribution to amino acid metabolism than the host itself, indicating a symbiotic relation ^47^. Similarly, in Pyropia haitanensis, co-cultivation with a Bacillus sp. did not only result in an increased growth rate, but also in a downregulation of genes related to the biosynthesis of several amino acids and other metabolites ^48^. As free amino acids are known to increase primary production of algae ^49,50^, it is hypothesised that the release of amino acids by bacteria serves as one mechanism through which they may facilitate algal growth.

### Large potential for vitamin B production by bacteria

Previous metagenomic studies in the red alga Pyropia haitanensis ^20^ and brown alga Nereocystis luetkeana ^21^ have highlighted the potential of the bacterial symbionts to produce vitamin B_12_ (cobalamin). Our data showed that the capacity of seaweed-associated bacteria to biosynthesize vitamins is widespread within the Ulva microbiome and extends beyond vitamin B12 (Fig. 1a). Vitamins are essential to a well-functioning central metabolism in both microbes and their hosts ^51,52^. Algae require a combination of different vitamins for growth, but are likely unable to synthesise some of these organic compounds themselves ^53–55^. A study by Croft et al. ^56^, for example, showed that more than half of the algae studied by them required an exogenous supply of vitamin B_12_ (cobalamin), 22% required vitamin B1 (thiamine), and 5% required vitamin B7 (biotin).

In Ulva species, the addition of vitamin B_12_ and particularly vitamin B_6_ (pyridoxal) to culture medium is necessary for growth and promotes nitrogen metabolism ^57,58^. This vitamin auxotrophy suggests that the host is dependent on the production by its microbial symbionts. Within our dataset, the capacity of bacteria to produce tetrahydrofolate (a derivative of vitamin B9; KEGG module M00126) was most widespread and was identified in 110 MAGs from varying families, including the Alteromonadaceae, Flavobacteriaceae, Granulosicoccaceae, Saprospiraceae, and Spirosomaceae (Fig. 1a). This vitamin was the only vitamin that was part of the core functions of the Ulva microbiome, being consistently present in all samples with a minimum relative abundance of 0.1%. KEGG modules related to the production of vitamin B_1_ (thiamine; M00127), vitamin B_2_ (riboflavin; M00125), vitamin B_5_ (pantothenate; M00119), vitamin B_6_ (pyridoxal; M00124), vitamin B_7_ (biotin; M00123), and vitamin B12 (cobalamin; M00122) were identified as well (Fig. 1a), although not as core functions. This could be attributed to either the absence of MAGs with the capacity to produce these vitamins in all samples or their presence in samples but not with high abundance.

### The Ulva microbiome is defined by its ability to degrade the host

Our metagenomic dataset provided a thorough overview of the functional profile of the Ulva-associated microbiome. Next, we aimed to determine the functions contributing to a “typical” Ulva microbiome, by contrasting taxa isolated from Ulva with those isolated from seawater. In total, we selected 152 MAGs from our metagenomic dataset, representing 33 different genera (Table S4). We then searched for publicly available genomes of bacteria from the same genera but isolated from seawater (71 genomes) (Table S4). Subsequently, we conducted a comparative analysis based on odds ratios to identify potential enrichments of specific KEGG Orthology (KO) terms or Carbohydrate-Active enZymes (CAZys) in bacteria from the same genus collected from Ulva versus seawater.

Our results suggest that a defining aspect of an Ulva microbiome is its ability to utilize and break down the host organism, shown by the significant enrichment of CAZys (incl. ulvan lyases, α-L-rhamnosidases, and rhamnogalacturonan α-L-rhamnohydrolase) that specifically target and break down Ulva’s polysaccharides and cell wall components (e.g., ulvan, iduronic acid, cellulose, xyloglucan) (Fig. 1b), as well as ABC transporters that facilitate the extracellular uptake of small monosaccharides like fructose (frcBCA; KEGG KO K10552, K10553 & K10554), α-glucoside (aglEFGK; K10232, K10233 & K10234), and rhamnose (rhaSPQT; K10559 & K10560) resulting from the degradation of the cell wall polysaccharides. It is clear that Ulva-associated bacteria do not only live on the algal tissue, they live of it as well.

### High taxonomic turnover and functional stability across salinity gradient

Subsequently, we examined the shifts in both the taxonomic and functional composition of Ulva-associated bacteria across a salinity gradient traversing four distinct salinity regions: spanning from the horohalinicum (5–8 PSU), through the mesohaline (8–18 PSU) and the polyhaline (18–30 PSU), to the euhaline (30–35 PSU) (Fig. 2a). The Atlantic–Baltic salinity gradient explained more of the observed variation in the taxonomic composition of Ulva-associated bacterial communities (p = 0.0001 , R^2^ = 0.70) than of the observed variation in the functional composition of the same communities (p = 0.0006, R^2^ = 0.17) (Fig. 2b, 2c). Pairwise comparisons, for example, showed that the taxonomic composition of Ulva-associated bacteria differed between all salinity regions (horohalinicum vs mesohaline vs polyhaline vs euhaline, p<0.01 for all comparisons; pairwise Adonis test). On the other hand, the functional gene profile of bacterial communities in the mesohaline, polyhaline, and euhaline were not significantly different from each other (p>0.05 for all comparisons, pairwise Adonis test), as only the horohalinicum differed from the two higher salinity regions (horohalinicum vs polyhaline, p=0.04; horohalinicum vs euhaline, p=0.04, pairwise Adonis test).

**Fig. 2.**
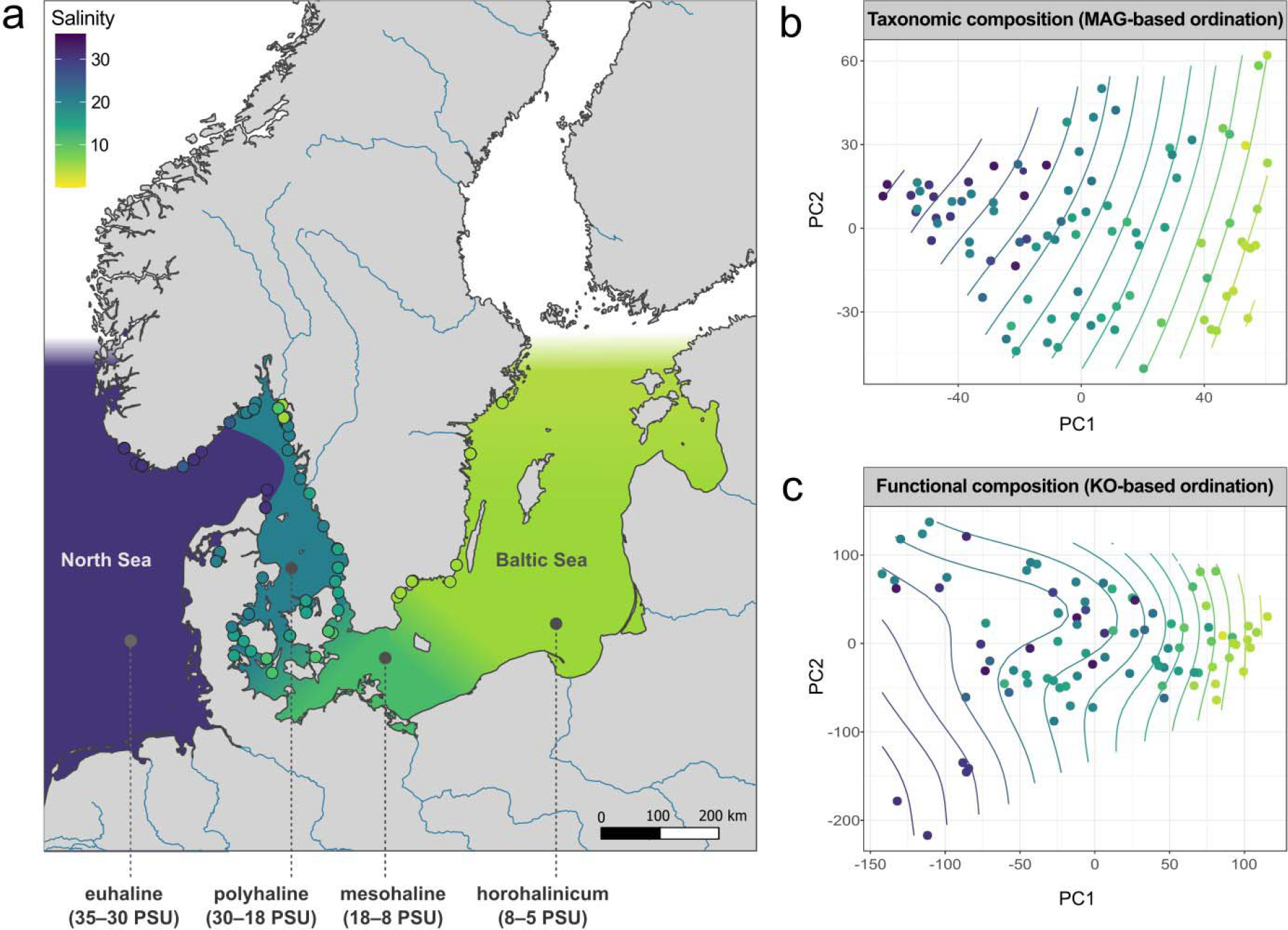
Taxonomic and functional patterns across salinity. (a) Geographic distribution of sampling sites (n=63) along the Atlantic–Baltic Sea gradient where a total of 91 Ulva sensu lato samples were collected. The four major salinity regions (horohalinicum, mesohaline, polyhaline, and euhaline) have been indicated on the map and rivers are projected in blue. PCA plots show the taxonomic (b) and functional (c) composition of Ulva-associated bacterial communities across salinity. The ordination in the taxonomic composition is MAG-based (metagenome-assembled genome) and the functional composition is KO-based (KEGG Orthologies). The contour lines (smooth surface lines) were fitted to the ordination plots based on the correlation with salinity. Colour represents salinity in all plots.

Nutrient concentrations, temperature and oxygen concentrations had little to no effect on taxonomic turnover (NOx, p = 0.04, R^2^=0.07; PO4, p = 0.08, R^2^=0.05; temperature, p = 0.01, R^2^=0.10; oxygen, p = 0.19, R^2^=0.04) and did not affect functional composition (NO_x_, p = 0.90, R^2^=0.002; PO_4_, p = 0.22, R^2^=0.03; temperature, p = 0.15, R^2^=0.04; oxygen, p = 0.84, R^2^=0.004).

Indicative of a high taxonomic turnover, 294 MAGs changed in relative abundance across the salinity gradient (p<0.01, LinDA linear regression) (Fig. 3a) (Table S2), of which 126 MAGs decreased with salinity and 168 MAGs increased with salinity. Several MAGs belonging to Dokdonia [MAG082, MAG518], Leucothrix [MAG022, MAG360], and Litorimonas [MAG193, MAG149] for example increased with salinity, while Alteraurantiacibacter [MAG014, MAG020, MAG578], Rubripirellula [MAG422], and Blastomonas [MAG197] decreased with salinity (Fig. 3b).

**Fig. 3.**
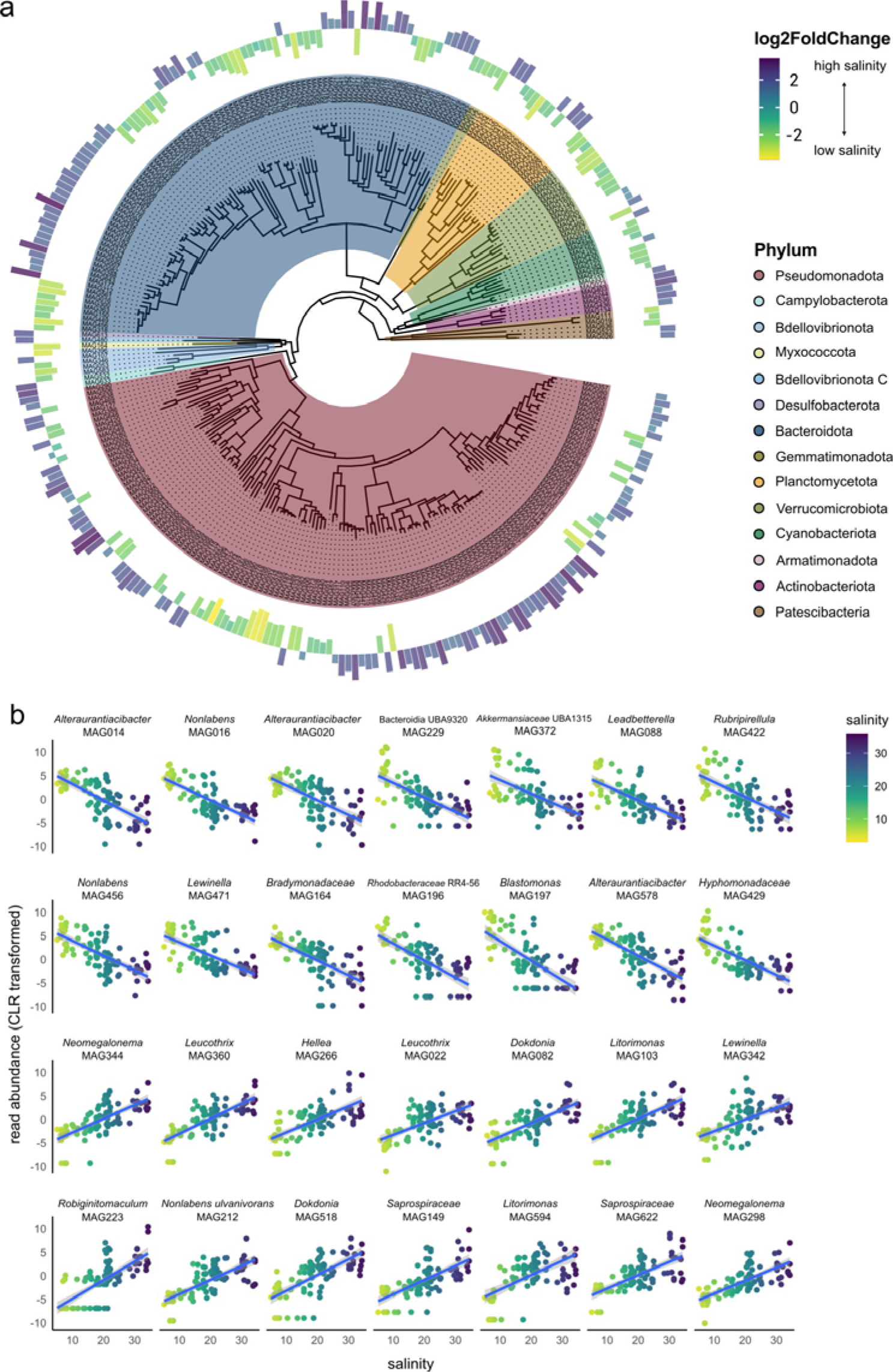
Abundance of metagenome-assembled genomes across salinity. (a) Phylogenetic tree of the 294 metagenome-assembled genomes (MAGs) that significantly differed (p<0.01) in relative abundance across the salinity gradient. The outer ring represents the log2FoldChange (LinDA linear regression model), with the MAGs in yellow-green that are enriched in low salinity and the MAGs that are enriched in high salinity in purple-blue. The colours in the phylogenetic tree represent the bacterial phyla (the legend is ordered in clockwise order of appearance). (b) Read abundance of the 28 most differentially abundant MAGs across salinity. The x-axis displays salinity (PSU) and the y-axis CLR-transformed read abundance. Colours represent salinity. Curves were fitted with a generalized linear model (GLM) using the R package ggplot2. Shaded areas represent the 0.95 confidence interval.

The taxonomic turnover was larger than the observed functional change, implying that multiple taxa across the salinity gradient were able to perform similar functions. Fig. 4 gives a complete overview of the presence or absence of KEGG modules in the metagenomic assembled genomes of 26 taxa that are characteristic to a specific salinity region (horohalinicum 5–8 PSU, mesohaline 8–18 PSU, polyhaline 18–30 PSU, or euhaline 30–36 PSU). Pantothenate (vitamin B_5_), for example, can be produced at low to medium salinity (5–18 PSU) by an unknown Saprospiraceae (MAG591), but at medium to high salinity (18–36 PSU) this function in the community was taken over by Robiginitomaculum and another unknown Saprospiraceae (MAG149) (Fig. 4). Similarly, proline biosynthesis at low salinity can be conducted by Alteraurantiacibacter and Rubripirellula (5–8 PSU), at medium salinity by Litorimonas A (8–18 PSU), and was gradually taken over by Neomegalonema (18–30 PSU), and Leucothrix and Litorimonas at high salinity (30–36 PSU). In addition, most MAGs shared a set of core genes that are necessary for essential functions, including nucleotide metabolism, fatty acid biosynthesis, energy metabolism (F-type ATPases, succinate dehydrogenases, etc.), and genes encoding structural modules (e.g., ribosomes, RNA polymerase, and DNA polymerase) (Fig. 4).

**Fig. 4.**
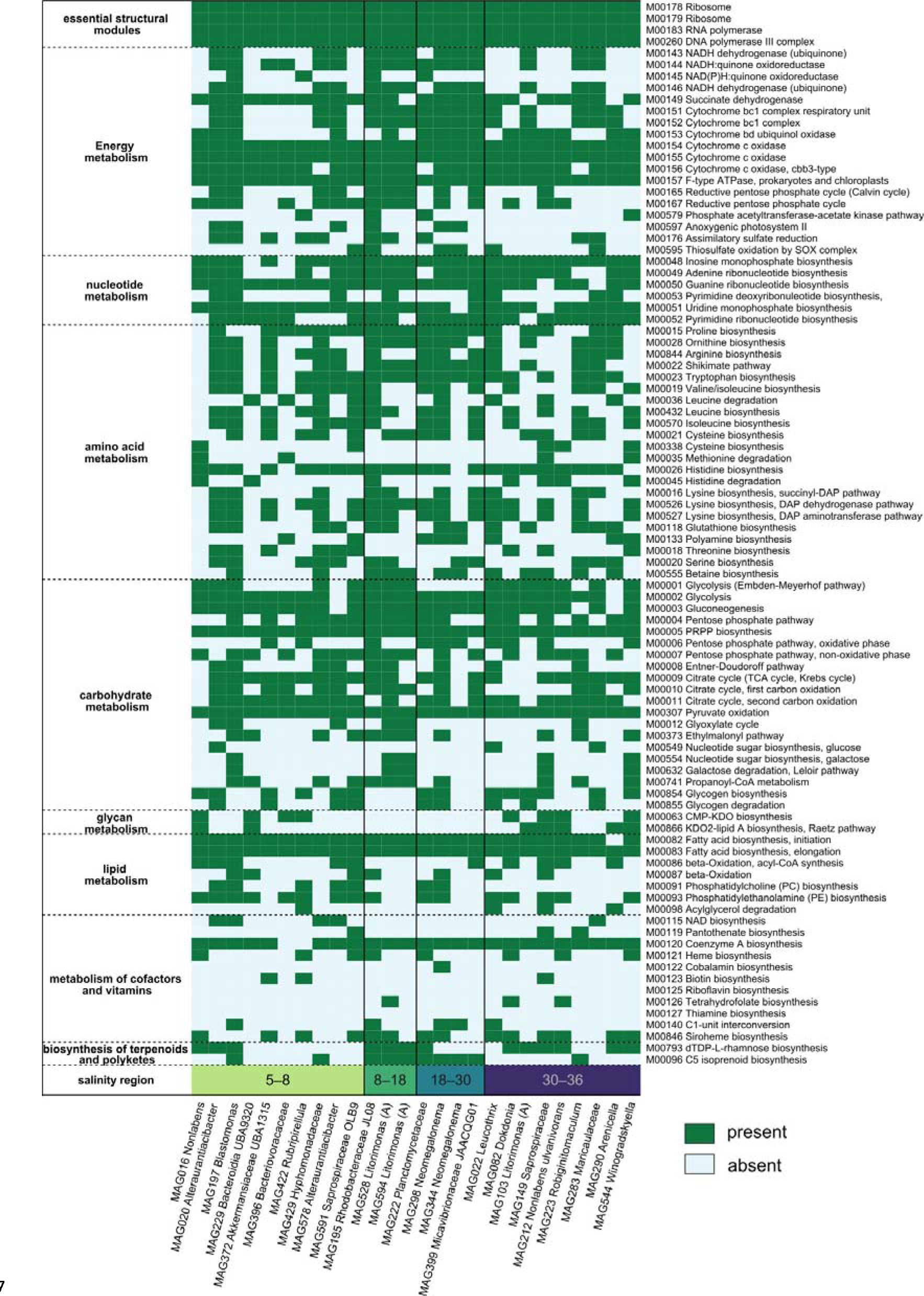
Taxonomic turnover and functional stability. Heatmap depicting the presence (dark-green) or absence (light-blue) of KEGG modules in 26 metagenome-assembled genomes (MAGs) that are characteristic to a specific salinity region (horohalinicum 5–8 PSU, mesohaline 8–18 PSU, polyhaline 18– 30 PSU, or euhaline 30–35 PSU).

### Osmoregulation drives functional variation

Despite lower functional turnover, we also identified 23 KEGG modules that differed in abundance across the salinity environment (p<0.05, LinDA linear regression) (Fig. 5). These modules are likely involved in the osmoregulation of bacterial cells, but may also affect the osmoregulation and acclimation potential of the host. One of the best-known strategies in bacteria and eukaryotes alike is the accumulation of low molecular weight compounds, such as sugars and amino acids, that act as osmoprotectants to maintain osmotic homeostasis and turgor pressure. Other strategies included the stabilization of cell membranes and mitigating oxidative stress.

**Fig. 5.**
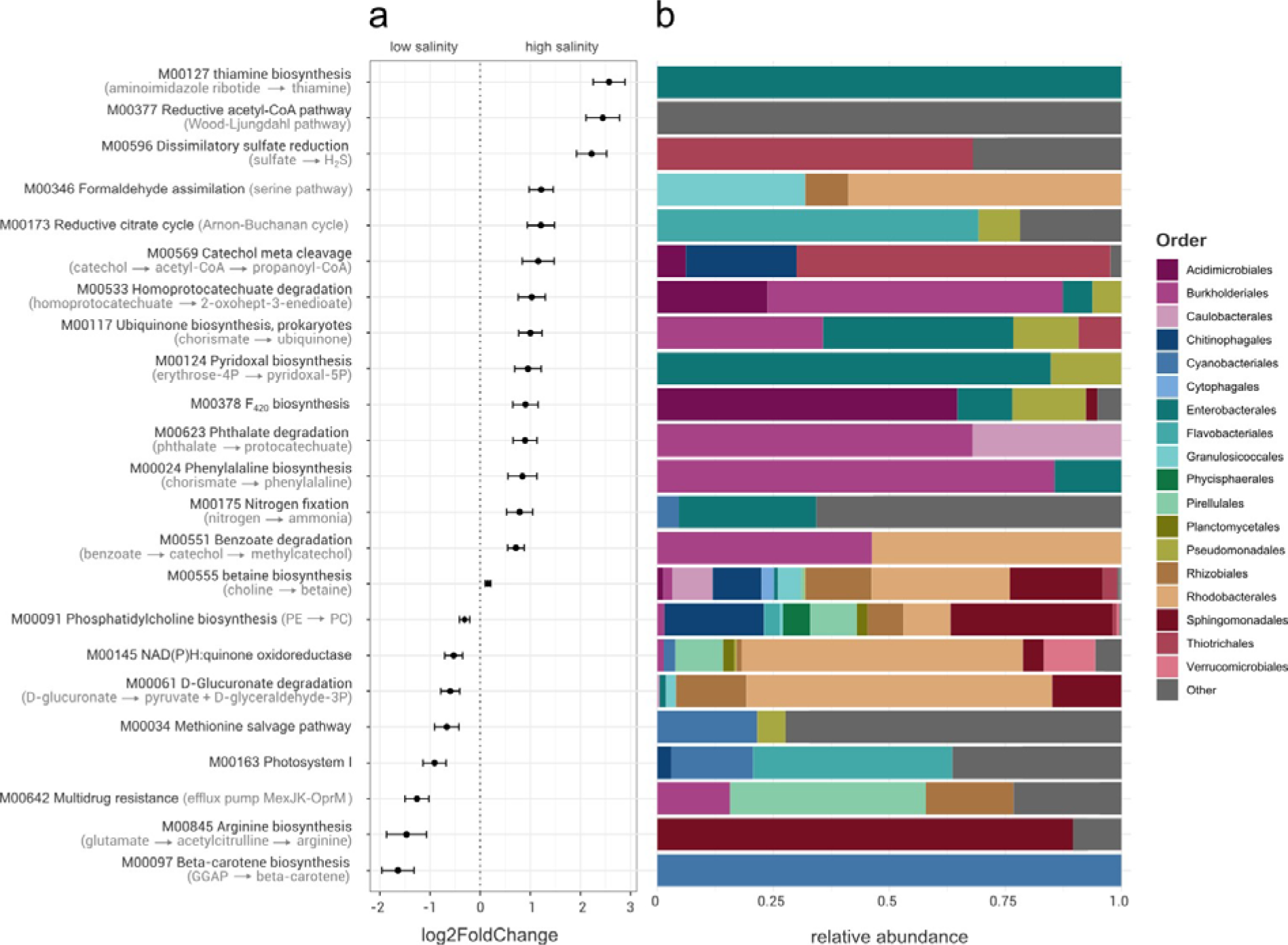
Functional changes in the Ulva-associated bacterial communities across salinity. (a) log2FoldChange values (± standard error) of the KEGG modules that significantly (p<0.05) differed in abundance with salinity. (b) Taxonomic composition (order level) of the metagenome-assembled genomes (MAGs) that harboured the identified KEGG modules. The relative abundance is based on read counts of the orders across the entire dataset.

Salinity-induced osmotic changes trigger the overproduction of reactive oxygen species (ROS), causing oxidative stress ^59^ and damaging membrane lipids, proteins, nucleic acids, and chloroplasts ^60^. Bacteria that were abundant in high salinity areas more often contained the genes needed to produce factor F420 (KEGG module M00378, p=0.0004), thiamine (vitamin B_1_; KEGG module M00127, p<0.0001) and pyridoxal-5P (the active form of B_6_; M00124, p=0.001). Several antioxidant mechanisms are F_420_-dependent ^61^ and cofactor F420 is known to help bacteria combat oxidative stress ^62^. In terrestrial plants, both vitamin B_1_ and B_6_ are known to alleviate salinity stress, for example in Arabidopsis thaliana ^63^ and in milk thistle ^64^, due to the stimulation of antioxidant enzyme activity and proline content. Similarly, gene expression of vitamin B_1_ is upregulated in phytoplankton during salt stress and oxidative stress is thought to function as a stress signalling molecule ^65,66^. In addition to protection from oxidative stress, pyridoxal-5P also increases photosynthetic pigment in wheat ^67^, and it facilitates growth through the reduction of ethylene accumulation that usually occurs under salinity stress in rice ^68^. As algae are likely not always able to produce their own vitamins ^53^, it is possible that thiamine and pyrodixal-5P produced by bacteria also aid the Ulva cells against oxidative radicals that are more prevalent under high salinity levels.

Bacteria that were abundant at high salinity also more often contained the genes needed to produce betaine (M00555, p = 0.03), an osmolyte that acts as one of the preferred compatible solutes in the majority of prokaryotes ^69,70^. It is known to accumulate in bacterial cells with increasing salinity ^71^, and betaine transporters are enriched in bacterial communities originating from marine habitats compared to freshwater environments ^72^. Although betaine plays a more important role in higher plants than in algae as osmoprotectant ^73^, a transcriptomic study showed that three choline dehydrogenase genes (involved in the conversion from choline to betaine) were upregulated in Ulva compressa during a recovery period after hyposaline stress ^74^. We also found an enrichment at high salinity of bacterial genes necessary for the production of the essential amino acid phenylalanine, which is known to accumulate in plants and algae under high salinity conditions ^75,76^ and increases salinity tolerance in maize ^77^. Genes associated with other well-known osmolytes and osmoprotectants, such as ectoine and trehalose, were comparatively less prevalent in our dataset. However, it is important to keep in mind that these findings are solely derived from gene content analyses and do not reflect gene expression patterns. Future transcriptomic studies could shed more light on the up- or downregulation of specific genes by Ulva and its associated bacteria in response to changing salinity conditions.

Another well-known strategy of bacteria to cope with cell turgor pressure is to alter the membrane composition through changes in fatty acids or phospholipids ^78^. The cell membrane separates the cell’s interior from the external environment and is therefore the first structure to encounter the effects of fluctuating salinity and osmotic stress. The disruption of membranes affects many processes such as transport of compounds, enzyme activities and signal transduction. It is therefore important to maintain the correct fluidity of the lipid bilayer ^79^. We found that many typical high-salinity MAGs were associated with the ability to synthesize ubiquinone (M00117, p=0.02), which agrees with the findings by Dupont et al. ^72^ on pelagic bacterial communities. Ubiquinone (also called coenzyme Q) is a membrane-stabilizing isoprenoid and the accumulation of this compound increases salt tolerance in bacteria, especially in the thin-walled gram-negative bacteria ^80,81^. Ubiquinone alters the physicochemical properties of the membrane by increasing the lipid packing and density. This results in reduced membrane permeability (i.e., a slower release of small solutes) and increases the strength of the membrane (i.e., resistance to cell rupture) ^82^. Conversely, we found that the module for phosphatidylcholine biosynthesis was enriched in low salinity conditions (M00091, p=0.002). Phosphatidylcholine (PC) is a membrane-forming phospholipid that is synthesized from Phosphatidylethanolamine (PE) and studies found that PE levels increased in salt-adapted cells ^83^. It has been noted before that while PE is the more common phospholipid in bacteria, gram-negative bacteria with high proportions of unsaturated fatty acids often contain additional PC to maintain stable bilayers ^84^. Indeed, in our dataset, the ability to convert PE to PC was mainly found in low salinity enriched Sphingomonadales (e.g., Alteraurantiacibacter, Erythrobacter) and Rhodobacterales (e.g., Jannaschia, Pseudorhodobacter). Membranes lacking PC are more fluid, have a higher permeability for small molecules and are more sensitive to osmotic changes ^85^. Both quinone and phosphatidylcholine seem to stabilize the membrane to withstand changes in turgor pressure and maintain osmotic balance.

### Bacteria-mediated acclimation

Our characterization of the metabolic functions of a typical Ulva-microbiome, highlights the large metabolic potential inherent to bacterial-algal symbiosis. The holobiont concept, which regards the seaweed host and its associated microbes as an integrated functional unit, is essential when studying the physiological response of seaweeds to environmental change. This work demonstrated that both the taxonomic and functional composition of Ulva-associated bacterial communities change across a 2,000 km salinity gradient. While Ulva-associated bacterial taxa displayed high taxonomic turnover across salinity, several Ulva species were able to colonize the entire gradient. This indicates that Ulva-bacteria have a smaller salinity-based niche than the host and are less tolerant to changes in salinity conditions (i.e. limited acclimation potential). The high turnover of microbial taxa is accompanied by functional redundancy, where guilds of taxa along the entire environmental gradient can perform crucial functions. These functions, including amino acid and vitamin B production, are potentially important to the seaweed host. Alongside functional redundancy, we identified distinct functional modules exhibiting enrichment in either low or high salinity areas. These modules are likely involved in mitigating oxidative stress, maintaining cellular osmotic homeostasis, and stabilizing cell membranes. Ulva depends on its microbiome for morphological development and growth — likewise, the forementioned bacterial acclimation mechanisms may play a role in host metabolism and acclimation. In light of bacteria-mediated acclimation, future laboratory experiments — involving the inoculation of seaweed cultures with targeted microbial communities — will be necessary to investigate whether bacteria can indeed facilitate the acclimation of Ulva species to changes in salinity.

## Methods

### Sample collection

Samples of Ulva sensu lato individuals (n=91) were collected during June–August 2020 in the Baltic Sea area (Table S1). Of each individual, a tissue sample was collected to molecularly identify the host species and a swab sample for microbiome analyses was generated by rubbing for 30 s on the tissue. Sterilized disposable gloves and sterilized equipment were used throughout the sampling procedure to minimize contamination.

All samples were stored in a portable freezer (-20 °C) until transferred to -80 °C in the laboratory.

In total, 63 sampling sites were visited along a salinity gradient in the Baltic Sea and adjacent areas such as the Kattegat, Skagerrak, and the eastern North Sea (Fig. 1a). The salinity ranged from 5.1 to 34.3 PSU and is presented in the figures in this study either on a continuous scale (0–35 PSU) or in salinity zones defined according to the Venice classification system (5–8 = horohalinicum, 8–18 = mesohaline, 18–30 = polyhaline, and 30–35 = euhaline) ^86^. In addition, water temperature (°C), oxygen levels (mg L^-1^), and nutrients concentrations (NO ^-^, NO ^-^, silicate, PO ^3-^ in µmol L^-1^) were measured at each site (Table S1).

Molecular identification, based on the tufA marker, confirmed the samples in our dataset represented seven different species: Blidingia minima (Nägeli ex Kützing) Kylin (n=8), Ulva compressa Linnaeus (n=10), Ulva fenestrata Postels & Ruprecht (n=8), Ulva intestinalis Linnaeus (n=29), Ulva lacinulata (Kützing) Wittrock (n=10), Ulva linza Linnaeus (n=20), and Ulva torta (Mertens) Trevisan (n=6) [see van der Loos et al. ^25^ and Steinhagen et al. ^23^ for detailed molecular methods and additional results concerning Ulva diversity in the Baltic region]. Throughout this study, “Ulva” refers to Ulva sensu lato (including Blidingia).

### DNA extraction and metagenomic sequencing

Total microbial DNA of the swab samples was extracted with the Qiagen DNeasy mini kit following the manufacturer’s protocol, with the addition of a bead beating step before lysis using zirconium oxide beads (RETCH Mixer mill MM400; 5 minutes at 30 Hz).

Quantity and quality of the DNA extracts were verified with Qubit (Life Technologies, Grand Island, USA) and NanoDrop (Thermo Scientific, Wilmington, USA). DNA extracts were sent to Novogene (Cambridge, United Kingdom) for library preparation and metagenomic sequencing on an Illumina NovaSeq 6000 (150 bp paired-end). A negative DNA extraction control and a positive control (ATCC microbial standard MSA-1002) were included. A total of 4,297,091,260 reads were generated (35,622,356–78,544,930 reads per sample). The sequences are archived at SRA (BioProject PRJNA1040445).

### Bioinformatics and statistical analyses

The metagenomic sequencing data was processed with the ATLAS Snakemake workflow^87^, which integrates quality control, assembly, genomic binning, and annotation. In short, quality control was performed using the BBTools suite ^88^. This includes removal of PCR duplicates and adapters, trimming and filtering of reads based on quality and length, and compressing the raw data files. Host sequences were removed based on an available Ulva reference genome (BioProject PRJEB25750) ^89^. The de novo metagenome assembly was done using MEGAHIT v1.0 ^90^. Next, metagenome-assembled genomes (MAGs) were predicted with binning tools MetaBAT 2 ^91^ and MaxBin 2.0 ^92^. Binning results were aggregated with DAS Tool ^93^. Quality assessment of the resulting MAGs was performed with CheckM ^94^. MAGs were dereplicated across samples with dRep ^95^, and taxonomic classification was performed with GTDB-Tk ^96^. Finally, genes were predicted with Prodigal ^97^, redundant genes were clustered with linclust ^98^, and annotated with eggNOG-mapper ^99^. This resulted in a classification of the genes following the Carbohydrate-Active EnZymes database (CAZy) ^31^ and Kyoto Encyclopedia of Genes and Genomes (KEGG) ^32,33^ databases.

The effect of salinity, NO_x_, and PO_4_ on bacterial community composition at both MAG (taxonomic profile) and KO (KEGG Orthology; functional profile) levels was assessed using the envfit function from the vegan package with 9999 permutations ^100^. Multivariate comparisons with 9999 permutations were subsequently conducted with pairwise Adonis among the different salinity zones ^101^. Functional differences across salinity were visualised with a PCA and smooth surface lines were fitted to the ordination with the ordisurf function (vegan package) based on the correlation with salinity ^100^. A LinDA linear regression was used to identify which MAGs and which KEGG modules (a set of genes with a specific reaction within a metabolic pathway) significantly changed across salinity and nutrient concentrations ^102^. P-values were Benjamini & Hochberg corrected. Given the compositional nature of the data, read abundance values were transformed with the centric log-ratios (CLR) prior to the analyses ^103^.

To gain a deeper understanding of what defines the metabolic potential of Ulva microbiomes, we compared the genomes of bacterial taxa isolated from Ulva to those isolated from seawater. In total, we selected 152 MAGs from our metagenomic dataset, representing 33 different genera. We then searched for publicly available genomes of bacteria from the same genera that were isolated from seawater (resulting in a set of 71 genomes) (Table S4). These originated from a variety of geographical locations and habitats (e.g., surface water, deep sea, hydrothermal systems, and oceanic gyres). Subsequently, we conducted a comparative analysis based on odds ratios to identify potential enrichments of specific KO terms or CAZy families within bacteria of the same genus collected from Ulva versus seawater. For each KO term and CAZy family, the odds ratio was calculated as the number of discordant genome pairs in favour of the Ulva (term/family present in the Ulva bacterial genome, but not present in the seawater bacterial genome) divided by the number of discordant pairs in favour of seawater (term/family present in the seawater bacterial genome, but not present in the Ulva bacterial genome). An offset of 0.5 was added to the number of discordant pairs to prevent dividing by zero. As multiple genomes were available per genus, pairs were randomly assigned 1000 times (permutations) and odds ratios were calculated for each permutation. The median odds ratio was retained. A term/family more frequent in Ulva bacterial genomes results in an odds ratio larger than one. The opposite results in an odds ratio smaller than one.

All statistical tests were performed in R ^104^ and data were visualised using the ggplot2^105^ and phyloseq ^106^ packages.

### Data availability statement

Raw whole-genome sequence reads and related metadata are deposited in the SRA (BioProject PRJNA1040445).

## Acknowledgements

The research leading to the results presented in this publication was carried out with infrastructure funded by the FWO PhD Fellowship fundamental research (3F020119), the EMBRC Belgium (FWO project I001621N), and the Formas-funded ‘A manual for the use of sustainable marine resources’ project (Grant no. 2022-00331). We would like to thank Samanta Hoffmann for her assistance during field work and Nadja Stärck for assistance with nutrient analyses. WS is funded through the FWO grant nr 1252821N.

## Author contributions

L.M.L.: Conceptualization, Methodology, Formal analysis, Data Curation, Writing - Original Draft, Visualization. S.ST.: Investigation, Writing, Funding acquisition - Review & Editing. W.S.: Resources, Writing - Review & Editing. F.W.: Writing - Review & Editing. S.D.: Methodology, Resources, Writing - Review & Editing. A.W.: Writing - Review & Editing, Supervision. O.D.C.: Writing - Review & Editing, Supervision, Funding acquisition.

